# Segmentation-free measurement of locomotor frequency in *Caenorhabditis elegans* using image invariants

**DOI:** 10.1101/2024.01.16.575892

**Authors:** Hongfei Ji, Dian Chen, Christopher Fang-Yen

## Abstract

An animal’s locomotor rate is an important indicator of its motility. In studies of the nematode *C. elegans*, assays of the frequency of body bending waves have often been used to discern the effects of mutations, drugs, or aging. Traditional manual methods for measuring locomotor frequency are low in throughput and subject to human error. Most current automated methods depend on image segmentation, which requires high image quality and is prone to errors. Here, we describe an algorithm for automated estimation of *C. elegans* locomotor frequency using image invariants, i.e., shape-based parameters that are independent of object translation, rotation, and scaling. For each video frame, the method calculates a combination of 8 Hu’s moment invariants and a set of Maximally Stable Extremal Regions (MSER) invariants. The algorithm then calculates the locomotor frequency by computing the autocorrelation of the time sequence of the invariant ensemble. Results of our method show excellent agreement with manual or segmentation-based results over a wide range of frequencies. We show that compared to a segmentation-based method that analyzes a worm’s shape and a method based on video covariance, our technique is more robust to low image quality and background noise. We demonstrate the system’s capabilities by testing the effects of serotonin and serotonin pathway mutations on *C. elegans* locomotor frequency.

## INTRODUCTION

Locomotion is an important parameter in behavioral genetics. Changes in motor behavior can reflect the effects of genetic mutations, modulations of neurochemical compounds, and variations in the physiological states (1). For the roundworm *C. elegans*, a variety of studies investigating changes or defects in its locomotor frequency have given insights into complex biological processes, including neurochemical modulation, neuromuscular coordination, and neural circuit development (2–8).

Current methods for quantifying locomotor frequency include manual scoring and machine vision approaches. Traditional manual scoring is performed by counting the number of body bending movements of worms placed in a solid or liquid environment (9). Manual measurements are technically simple but low in throughput and subject to observer error.

To address these limitations, progress has been made in machine vision systems for automating the analysis of *C. elegans* locomotor movements. Most automated methods rely on analyses of animal morphometry. In these methods, each image undergoes filtering and thresholding to yield a binary image, from which features such as centroid position, body centerline, curvature, and posture can be determined. Extracting these features from image sequences enables further determination of behavioral dynamics, including speed, amplitude, reorientations, and locomotor frequency (7, 10–14). These methods are highly susceptible to recording conditions and are vulnerable to image noise, low image quality, and behavioral fluctuations. In addition, these methods often require some degree of manual intervention, such as for identifying key points (head and tail) and performing additional noise reduction steps (5, 10).

An alternative automated strategy computes the covariance matrix of video frames, inferring locomotor frequency by measuring the time intervals between frames that exhibit statistical similarity (15). This method is capable of automatically measuring worm locomotor frequency without using morphometry. However, this direct covariance between video frames is degraded by translation or rotation of the animal over time.

Here, we introduce a method we call Imaginera (Image Invariant Ensemble for Rhythmicity Analysis), which uses an image invariant ensemble to conduct rhythmicity analysis in freely behaving *C. elegans*. Our method extracts several affine-invariant image features from video frames and computes the autocorrelation of the time sequence of the invariant ensemble to infer frequency. We evaluated the robustness and speed of our method by testing its performance in analyzing videos of moving animals under varied spatiotemporal resolutions or in the presence of background noise and comparing outcomes to those of other methods. We applied our technique to a multi-well experimental setup to analyze the locomotion of hundreds of wild-type and mutant individuals under controlled environmental conditions. These results show that Imaginera is a robust, high throughput method for measurement of *C. elegans* locomotor frequency.

## RESULTS

### mplementation and performance of automated frequency tracking

Our analysis consists of three modules, each carrying out a sequence of image analysis operations on each video frame. The first module provides an initial background reduction of a video frame by subtracting the video background derived from the first principal component of the original video (15) (Fig. 1A).

**Figure 1.**
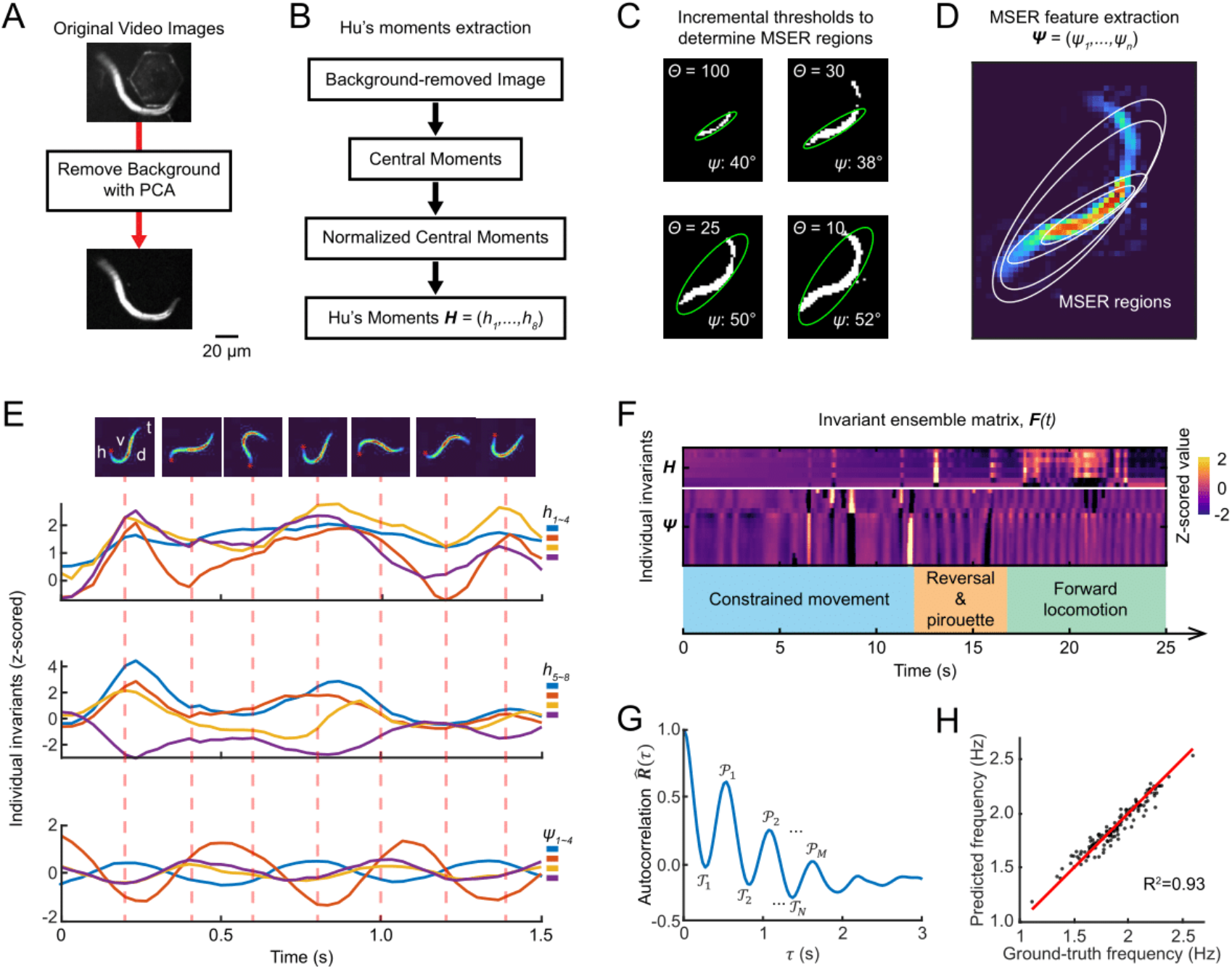
Implementation and performance of Imaginera. (A) Video images undergo background subtraction. (B) Calculation of Hu’s moment invariants from a background-removed video image. Hu’s moment invariants, ***H*** = (*h*_1_, *h*_2_, …, *h*_3_), are a series of 8 image moments derived from the normalized central image moments of order from 0 to 3. (C) The MSER detector incrementally thresholds a background-removed video image through the intensity range of the image to determine MSER regions. Each MSER region can be represented by an ellipse with identified orientation angle (*ψ*). (D) Orientation angles (***Ψ***) of the detected MSER regions. (E) Invariants from Hu’s moments and MSER regions are extracted from a sequence of video frames (representative frames shown on *Top*). Curves are representative examples of Hu’s moments 1-4 (*Panel 1*), Hu’s moments 5-8 (*Panel 2*), and orientation angles (*Panel 3*) of 4 MSER regions. (F) Invariants over time are combined to form the invariant ensemble matrix (***F***(*t*)). Colored rectangles indicate behavioral patterns. Constrained movement was observed from a worm confined by a microfluidic channel (5). (G) Autocorrelation of the invariant ensemble. *𝒫*_1_, *𝒫*_2_, …, *𝒫*_*M*_ denote local maxima and *𝒯*_1_, *𝒯*_2_, …, *𝒯*_*N*_ denote local minima. (H) Comparison between the frequencies calculated by the automated method vs. manual observation of recorded videos.

The subsequent module calculates a set of 8 Hu’s moment invariants from each background-subtracted frame (16) (Fig. 1B; see *Methods*). Each Hu’s moment invariant constitutes a specific weighted average (moment) of image pixel intensities or their positive-integer powers. These moments are invariant with respect to image translation, scale, and rotation (16).

The third module, termed the MSER (Maximally Stable Extremal Regions) detector, extracts a series of co-variant regions from an image, which are a sequence of stable connected components from certain gray-level sets of the image (17) (see *Methods*). This operation is executed by first sorting all pixels by gray value, then adding pixels incrementally to each connected component as the gray value threshold decreases (Fig. 1C; method valid for dark-field images). While monitoring the pixel area, regions that display minimal variation with respect to the threshold are designated Maximally Stable Extremal Regions, or MSERs. Elliptical frames are then marked by fitting minimal circumscribed ellipses to the regions, whence orientation angles (***Ψ***) are computed as feature invariants (Figs. 1 C and D; see *Methods*).

Lastly, the software standardizes the time sequences of the invariants extracted from the previous modules using z-score normalization (Fig. 1F) and computes the autocorrelation of the invariant ensemble (Fig. 1G; see *Methods*). Displayed within the autocorrelation curve is an alternating series of peaks (*𝒫*_*M*_) and troughs (*𝒯*_*N*_) reflecting periodicity of the object in the video (Fig. 1G). The frequency value is computed as the reciprocal of the mean of all intervals between consecutive peaks and troughs within the autocorrelation (Fig. 1G; see *Methods*).

To validate the accuracy of Imaginera in quantifying animals’ moving frequency, we acquired a testing dataset consisting of 153 videos of individual freely swimming worms, each at least 10 s in duration, recorded at 30 frames per second (fps) and a 6.7 µm pixel resolution (see *Methods*). We compared automated frequency measurements from Imaginera with ground-truth results from manual scoring within these videos and found an R-squared value of 0.93 (Fig. 1H). We found that the accuracy of the method decreased if any one of the three modules was disabled. Specifically, the R^2^ value dropped to 0.45, 0.71, and 0.75 when we disabled the background subtraction module, Hu’s invariants module, and MSER invariants module, respectively. These results show that our method, Imaginera, relies on its three modulates to provide an accurate estimate of *C. elegans* locomotor frequency.

### Performance under spatial or temporal downsampling

The spatial and temporal resolution of behavioral recordings can be limited by several factors including equipment constraints, data storage constraints, field of view, number of animals being imaged, and computational time. We investigated the extent to which the software could estimate frequency with imaging at lower spatial and/or temporal resolution. We downsampled our testing dataset in time and space to emulate image sequences with lower frame rates or lower pixel resolution (Figs. 2 A and B).

**Figure 2.**
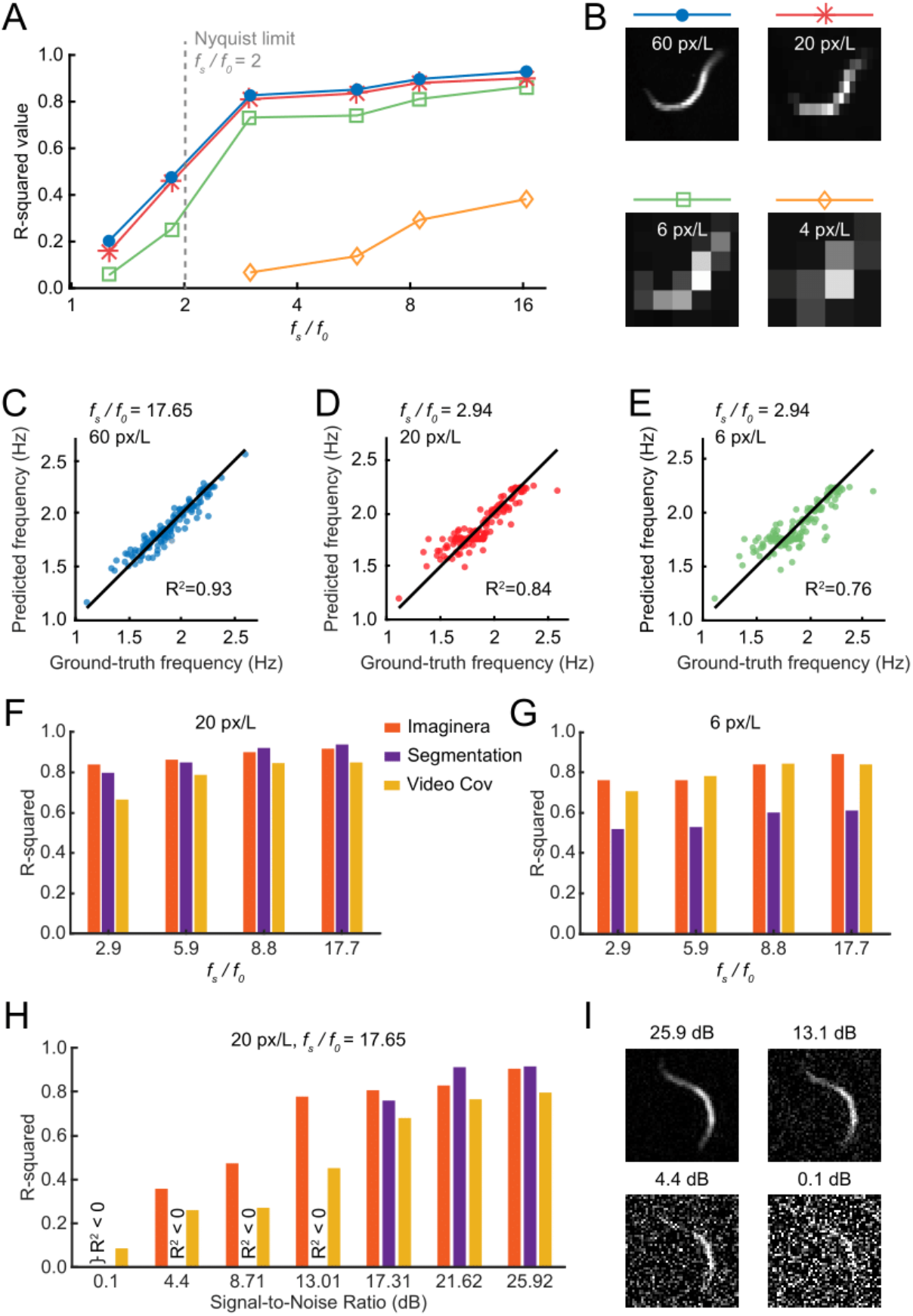
Robustness analysis of Imaginera. (A) Accuracy of the Imaginera method in measuring frequency under various recording conditions, emulated by computationally downsampling the spatiotemporal resolutions of the original videos. R^2^ shown as a function of the ratio of the video acquisition frequency (*f*_*s*_) and average locomotor frequency (*f*_0_ ≈ 1.7 *Hz*). Line colors indicate different spatial resolutions denoted by the number of pixels to resolve an average worm length *L*. The gray dashed line indicates the Nyquist limit for the average frequency. (B) Worm images under different spatial resolutions. (C-E) Comparison between results of the Imaginera method and manual observation for measuring locomotor frequency under indicated spatiotemporal resolutions. (F and G) Comparison of frequency measurement accuracy between methods using Imaginera, segmentation, and video covariance, under indicated spatiotemporal resolutions. (H) Comparison between these methods in measuring moving frequency from noisy videos under varying signal-to-noise ratio. (I) Representative noisy worm images with varied signal-to-noise ratios.

We first investigated the effect of changing temporal resolution while keeping spatial resolution fixed. We found that accuracy exhibited a mild decline as the temporal resolution decreased within a range in which the acquisition frequency (*f*_*s*_) was at least triple the animals’ average locomotor frequency (*f*_0_ ≈ 1.7 *Hz* and *f*_*s*_/*f*_0_ ≥ 2.94) (Fig. 2A). The accuracy plummeted when temporal resolution was around or below the Nyquist-limit critical point, twice the average locomotor frequency of the animal (*f*_0_ ≈ 1.7 *Hz*) (Fig. 2A).

Next, we asked how well the algorithm worked with lowered spatial resolution (Fig. 2A), keeping temporal resolution constant. The accuracy of frequency measurements remained stable as the spatial resolution reduced from 60 to 6 pixels per worm length (*L* ≈ 1 *mm*) (Fig. 2B). The accuracy dropped markedly when the spatial resolution decreased to 4 pixels per worm length.

We used Imaginera to analyze the testing dataset under different combinations of temporal and spatial resolutions. The algorithm showed excellent agreement with ground truth data (R^2^ = 0.93) when analyzing videos with a 30 fps temporal resolution (*f*_*s*_/*f*_0_ = 17.65) and a 6.7 μm spatial resolution (60 pixels per worm length) (Fig. 2C). The performance remained passable under relatively poor recording conditions where temporal resolution was 5 fps (*f*_*s*_/*f*_0_ = 2.94) and spatial resolution was 20 μm (20 pixels per worm length) or even 67 μm (6 pixels per worm length), yielding R-squared values of 0.84 and 0.76, respectively.

### Performance comparison with alternative methods

We compared the performance of the Imaginera software with two other methods for measurement of frequency: a widely used approach based on a segmentation-based algorithm to analyze the worm morphometric skeleton (10), and a non-morphometric method that computes the covariance matrix of video frames (15).

We used Imaginera and the two alternative methods to analyze the testing dataset across various spatiotemporal resolutions. When analyzing mildly pixelated videos (20 pixels per worm length), Imaginera delivered accuracies similar to or slightly better than the segmentation method and the video covariance method under varied temporal resolutions (Fig. 2F). When analyzing videos that are more coarsely pixelated (6 pixels per worm length), both Imaginera and the video covariance method noticeably outperformed the segmentation method (Fig. 2G).

To assess the extent to which Imaginera is robust to background noise, we computationally added Gaussian white noise to the testing dataset (see Fig. 2I), and tested the three methods on the noisy datasets with varied signal-to-noise ratio (SNR). When analyzing videos with SNR ≥ 17.31 dB, all three methods yielded similar accuracy (R^2^ ≥ 0.76), with Imaginera and segmentation-based methods performing consistently more accurate than the video covariance method (Fig. 2H). When SNR < 14 dB, worm images became noticeably noisy (Fig. 2I), and accuracy of the three methods further decreased, of which Imaginera exhibited the most robustness whereas the segmentation method failed due to susceptibility to noise (Fig. 2H).

To assess computational efficiency, we measured the time each method took to analyze 10,000 video frames (120 x 160-pixel resolution) and found that Imaginera executed analyses under 40 s, approximately 3 times faster than the segmentation-based method. While the implementations of these algorithms have yet to be optimized for speed, these results suggest that our method may be faster than the traditional segmentation method.

### Automated frequency analysis facilitates large-screen behavioral assays

Having demonstrated the accuracy and robustness of Imaginera under computationally emulated recording conditions (Fig. 2A), we evaluated its proficiency in practical applications by applying it to a multi-array imaging setup intended for large-scale phenotypic assays (Ji et al., *in preparation*).

During data acquisition, we employed a standard 96-well plate, with individual animals situated in wells filled with liquid NGM under defined external conditions. The imaging setup captured details at approximately one-sixtieth the length of an adult *C. elegans*, with a pixel resolution of 18 µm for a field of view that included all 96 wells (Figs. 3 A and B). In each experiment, up to 96 1-day adult animals were monitored for a duration of 5 hr. Every 2 minutes during this period, a 20-s-long image sequence was captured at 5 fps. We used Imaginera to analyze the movements of each animal in a well during each 20 s recording window.

**Figure 3.**
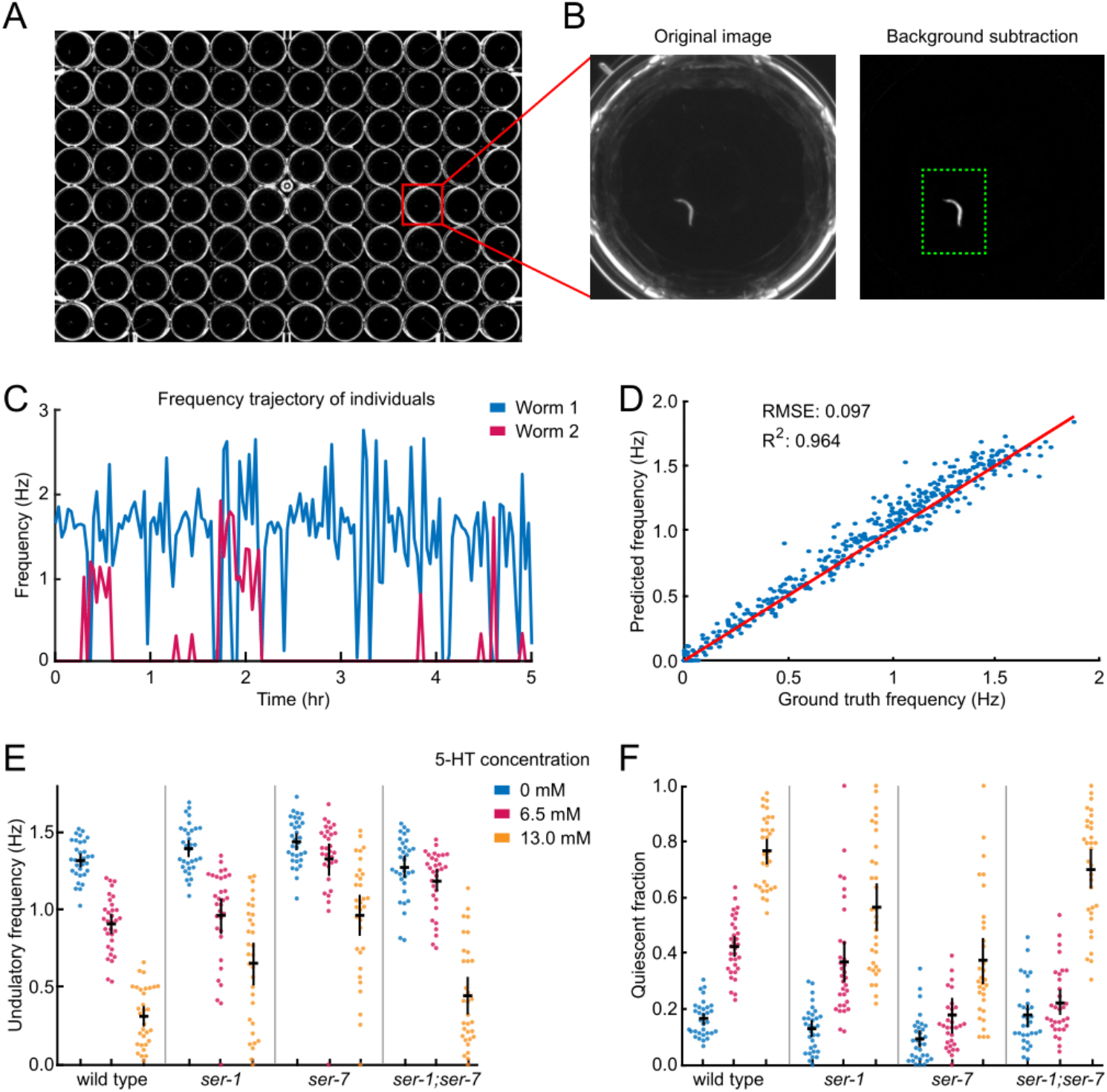
Automated frequency analysis facilitates large-screen behavioral assays. (A) Worms in a 96-well assay plate under dark field illumination (FOV 108 mm x 73 mm). (B) Individual wells are cropped and background-subtracted before analysis (FOV 6 mm x 6 mm). (C) Locomotor frequency dynamics of two individual animals calculated from a 5 h video sequence by the Imaginera method. (D) Locomotor frequency values evaluated by human observation (ground truth) compared with those from automated frequency assessment. (E) Locomotor frequency of wild-type and mutant individuals. (F) Quiescent fraction of wild-type and mutant individuals. In E and F, each point represents an animal, point colors indicate groups in the respective serotonin conditions, black bars represent population mean ± SEM.

The biogenic amine serotonin is an important neuromodulator that regulates *C. elegans* behavior in response to changing environmental cues (18). Exogenous serotonin at sufficiently high concentrations can cause inhibition of locomotion (19, 20). We tracked the frequency dynamics of wild-type and mutant *C. elegans* individuals in liquids with controlled serotonin (5-HT) concentrations. The mutant strains under examination included *ser-1, ser-7*, and *ser-1; ser-7* double mutants, each carrying loss-of-function mutations in the G protein-coupled metabotropic receptors SER-1 and/or SER-7.

We computed each animal’s level of behavioral quiescence by calculating the fraction of time the frequency equaled zero (Figs. 3 E and F). Generally, both wild-type and mutant animals exhibited normal locomotor frequency and low quiescence under serotonin-free conditions (Figs. 3 E and F). With the incremental addition of exogenous serotonin, wild-type animals exhibited a decline in the average locomotor frequency and increased quiescence (Figs. 3 E and F). We found that *ser-1* and *ser-7* mutants exhibited milder serotonin-induced effects on locomotor frequency and quiescent fraction, while *ser-1; ser-7* double mutants showed frequency slowing responses or quiescence to an intermediate level of exogenous serotonin and stronger responses to a high level of serotonin (Figs. 3 E and F). Our behavioral data align with previously reported results for each strain regarding the effects of exogenous serotonin on *C. elegans* locomotion (3, 19, 21–24).

We evaluated the accuracy of our method in this multi-well setup by comparing automated measurements to manual counts of head bends in 480 video clips (each 6 s long) of individual wild-type worms across all 96 wells. Our method demonstrated excellent agreement with results from manual counting (Fig. 3D).

These results show that our method is suitable for measuring frequency in 96 animals simultaneously.

## DISCUSSION

The Imaginera software provides a measurement of frequency in the locomotion of *C. elegans*, incorporating both non-morphometric invariants drawn from image profiles via Hu’s moments and morphometric invariants derived from object conformations via MSER regions. Our method thus complements other behavioral output metrics utilized in large-scale phenotypic assays, such as pixel differences between successive frames (2, 25).

We found that Imaginera has several advantages over existent morphometric and non-morphometric approaches. First, our method is more robust to errors that could arise from inaccurate separation of the worm from its background or from the incorrect abstraction of morphometric parameters. Second, our method minimizes the effect of noise introduced by sensitivity to translation and rotation in previous covariance-based, making Imaginera more accurate and robust when analyzing pixelated or noisy videos. Finally, because a threshold-based peak finding algorithm on the autocorrelation is implemented in Imaginera to determine frequency, it allows Imaginera to define weakly periodic or irregular motion of worms and assign zero frequency to these cases.

A key requirement for utilizing Imaginera is that worms must not overlap, as the movement of multiple worms in the same field of view complicates obtaining precise measurements from the autocorrelation of image invariants. To simultaneously measure movement of multiple worms, we have implemented this method for videos of 96 worms individually arrayed in a multi-well plate. Deployed in this way, we demonstrate the software’s high applicability in automating large-scale chemical and genetic screens for effects on movement.

In this study, we applied Imaginera for analyzing *C. elegans* locomotor rhythmicity; however, since Imaginera is established based on pixel intensities and does not involve any prior knowledge or constraint for worm’s morphology and movement, its utility could be adaptable to broader image analysis contexts that require quantifications of periodic behaviors or phenomena. This includes locomotor behavior in other animals like *Drosophila* larvae and zebrafish (26, 27), as well as other rhythmic dynamics like respiratory rhythms (28), pharyngeal pumping (29), microbial oscillations, and flight wingbeat (30). We anticipate that our method will aid in identifying subtle behavioral phenotypes and facilitating automated studies of periodic dynamics.

## METHODS

### C. elegans strains and maintenance

*C. elegans* strains were cultivated on *Escherichia coli* strain OP50 at 20 °C following standard methods (31). The strains used in this study include wild-type strain (Bristol N2), *egl-1(n487), ser-1(ok345), ser-7(tm1325)*, and *ser-7(tm1325); ser-1(ok345)* mutant strains, all obtained from the *Caenorhabditis* Genetics Center. All experiments were conducted with 1-day-old adult worms. To synchronize worm populations, we used a timed egg lay approach (32).

### Behavioral data acquisition

To capture the locomotion of freely moving worms, we placed animals in NGM buffer within a chamber formed by a glass slide and a coverslip, separated by 125-µm-thick polyester shims (McMaster-Carr 9513K42). This NGM buffer mirrors the composition of solid NGM, excluding agar, peptone, and cholesterol (33). The NGM buffer was supplemented with 0.1% (by weight) bovine serum albumin (BSA) to deter worms from sticking to chamber walls.

Behavioral footage of each worm within the field of view was recorded for approximately 1 min using a 5-megapixel CMOS camera (DMK33GP031, The Imaging Source, Inc.) paired with a C-mount lens (Nippon Kogaku NIKKOR-H; effective focal length 28 mm). Video sequences, captured at 30 fps with a 6.7-µm pixel resolution, were acquired using IC Capture software (The Imaging Source) under dark-field illumination provided by a red LED ring (outer diameter 80 mm; Qasim).

### Multi-well plate assays

For multi-worm imaging, we picked a single worm into each well of a standard 96-well microtiter assay plate (Corning 3795, round bottom), containing 60 µL of NGM buffer or NGM buffer containing serotonin.

The assay plate was positioned on a stage inside a custom-built imaging apparatus. Dark-field lighting was provided by an LED light ring (outer diameter 12 inches; Sunpak) above the platform. Videos were recorded with a CMOS camera (DMK 33UX183, 20 megapixels, The Imaging Source) coupled with a C-mount lens (HF12.5SA-1, Fujinon), recording at 5 fps for a duration of 20 s using IC Capture software (The Imaging Source). The behavioral assay involved obtaining a sequence of these 20 s videos, taken every 2 minutes until a cumulative recording time of 5 h was reached. The system’s field of view was calibrated to encompass all 96 wells on the plate, yielding a pixel resolution of 18 µm.

When analyzing multi-well videos, the Imaginera method is applied to the entire imaged well, and the positions of all wells in the videos are specified in a GUI designed in the software before the computational analysis by Imaginera.

### Mathematical formulae of Hu’s moment invariants

The Hu’s image invariants (16) with respect to translation, scale, and rotation are defined as:

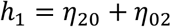

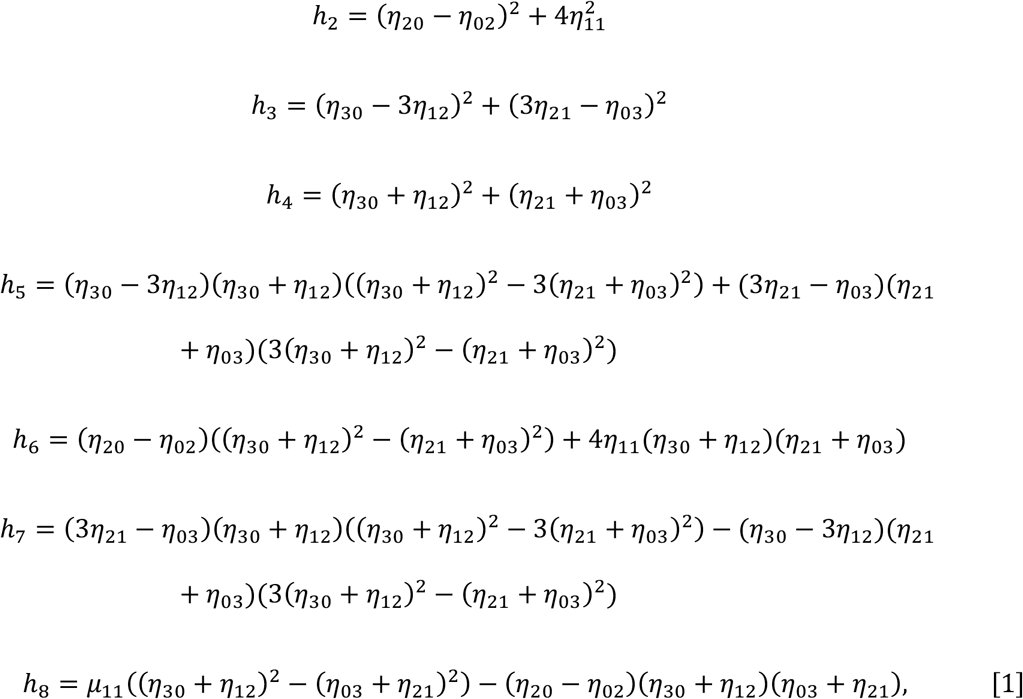

where 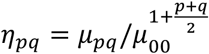 is the scaled central moment of order (*p* + *q*) and 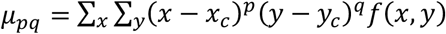 is the central moment of image intensity *f*(*x, y*) with the image centroid defined as 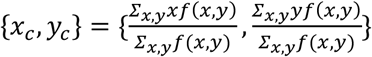.

Out of the 8 Hu’s moments, *h*_1_ through *h*_6_ are reflection-symmetric, meaning they remain unchanged with respect to image reflection, whereas *h*_7_ and *h*_8_ are reflection-antisymmetric, in that they invert their signs when the image is reflected.

### Extraction of MSER regions and features

The Maximally Stable Extremal Regions (MSER) are characterized by connected pixel components that depicted areas with a consistent intensity over a range of pixel values in the image (17). We constructed the minimal circumscribed ellipses that enclosed these regions, with the largest ellipse covering the entire worm and the smallest one covering the worm’s brightest segment. For each video frame, we calculated the orientations of the ellipses, resulting in a sequence of orientation values (termed feature invariants).

Given that the MSER method operates based on image intensity levels, the count of extracted ellipses may vary between frames due to variations in pixel intensity. To ensure uniformity in the number of feature invariants (*N* by default) across all frames, we computed *N* orientation values represented as ***Ψ*** = (*ψ*_1_, *ψ*_2_, …, *ψ*_*N*_) in radians for every frame, using bilinear interpolation and extrapolation methods with respect to *N* evenly spaced query values of connected component areas (in pixels) defined between predefined maximum and minimum values. To achieve rotational invariance in orientation values, the ***Ψ*** vector was recentered by deducting its circular mean, 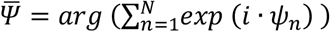 leading to the adjusted value 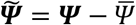.

### Ensemble feature invariants for determining locomotor frequency

We define an ensemble vector combining the 8 Hu’s Moment invariants (***H***) with the *N* MSER-derived invariants 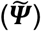, forming a vector with *P* = 8 + *N* elements:

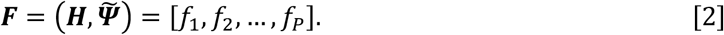

Next, we create an invariant ensemble matrix ***F***(*t*) that includes the time dependence of the vector sequence:

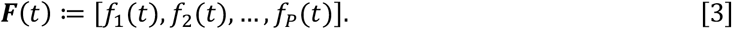

Each component variable *f*_*i*_(*t*) was measured on different scales and with widely different ranges in magnitude. We thus standardized them via z-score normalization:

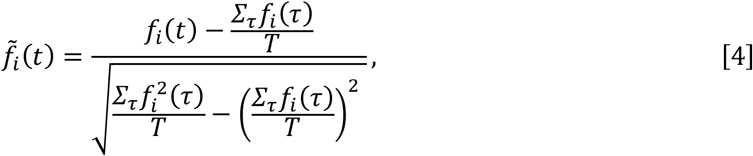

where *T* denotes the number of video frames.

Following the standardization of the invariant ensemble time sequence, we computed its autocorrelation 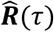, defined by:

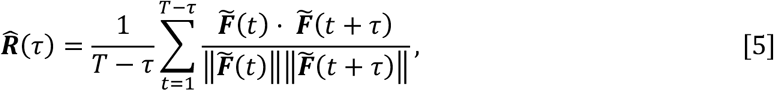

where 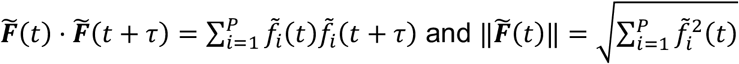.

We used the following peak-finding method to determine the locomotor period: 1. In 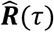, identify local maxima (peaks, including the one at *τ* = 0), {*𝒫*_1_, *𝒫*_2_, …, *𝒫*_*M*_}, and local minima (troughs), {*𝒯*_1_, *𝒯*_2_, …, *𝒯*_*n*_}. 2. Combine the peak list and the trough list, and reorder the joint list ascendingly based on the extrema’s *τ* values, which gives us {*ℰ*_1_, *ℰ*_2_, *ℰ*_3_, *ℰ*_4_, …, *ℰ*_*m*+*n*_} where *ℰ*_*2k−1*_ = *𝒫*_*B*_ and *ℰ*_2*k*_ = *𝒯*_*k*_. Here all the peaks and troughs must alternate in the joint list based on the Intermediate Value Theorem. 3. In the list {*ℰ*_1_, *ℰ*_2_, *ℰ*_3_, *ℰ*_4_, …, *ℰ*_*m*+*n*_}, compute all the identified local extrema’s topographic prominence, a measure of how much the peak or trough stands out relative to other peaks or troughs (34). 4. From the list *E* = {*ℰ*_1_, *ℰ*_2_, *ℰ*_3_, *ℰ*_4_, …, *ℰ*_*m*+*n*_}, find the family *P* of the least number of subsets of *E* that meet the following criteria: (a) The family *P* does not contain empty set or any set with only one element; (b) If the family *P* is not empty, each set in the family *P* does not contain any element with the prominence value smaller than 0.14; (c) If the family *P* is not empty, each set in the family *P* must contain at least two elements with their subscripts being adjacent integers; (d) If the family *P* is not empty, the intersection of any two sets is empty; (e) The union of all sets in *P* is equal to *E* − *X*, where *X* is the set containing all elements with their prominence values smaller than 0.14. 5. After identifying the family *P*, the worm’s locomotor frequency in the targeted video is defined as: (a) If the family *P* is empty, *f*_*worm*_ is defined to be 0; (b) If the family *P* is not empty, we define *P* = {*p*_1_, *p*_2_, …, *p*_*l*_} where 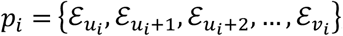, *i* = 1,2,3, …, *l*, and *f*_*worm*_ is calculated as 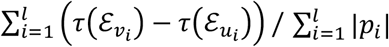, where | · | denotes the cardinality of the set.

### Segmentation-based approach for frequency analysis

In order to compare with our method, we used image thresholding and segmentation to extract the morphometric skeleton of worm images from videos (10, 12, 35). For each video frame, we filtered and thresholded the image to give a binary image that identifies the boundary of the worm. The head and tail of the worm were identified as the points of local maximal convex curvature of the worm boundary. A worm centerline extending from the head to the tail was calculated as points equidistant to the two sides of the worm boundary (7). and identified the worm’s boundary and centerline from the binary image. Next, we smoothed the worm’s centerline using a cubic spline fit and computed the curvature of the body centerline as the dot product between the unit normal vector to the centerline and the derivative of the unit tangent vector to the centerline with respect to the body coordinate.

To infer the animal’s movement frequency from the curvature dynamics, we defined anterior curvature as the average of the whole-body curvature across body coordinates 0.1-0.3 (fractional distance from the head). This specific range was chosen to exclude the high-frequency movements of the animal’s anterior tip (12). Lastly, we determined the local extrema from the anterior curvature and the frequency during the examined video period was defined as the product of the acquisition frequency and the reciprocal of the mean of all peak intervals in the anterior curvature.

## DATA AVAILABILITY

Strains are available upon request. The software and sample image data are available at https://github.com/cfangyen/Imaginera.

## ACKNOWLEDGMENTS

We thank Niels Ringstad, Yen-Chih Chen, and Zihao (John) Li for providing reagents and helpful discussions. Some strains were provided by the *C. elegans* Genetics Center, funded by the NIH Office of Research Infrastructure Programs (P40 OD010440). This work was supported by the National Institutes of Health (R01DA056358).

